# Synthetic lipid-containing scaffolds enhance production by co-localizing enzymes

**DOI:** 10.1101/052035

**Authors:** Cameron Myhrvold, Jessica K. Polka, Pamela A. Silver

**Author notes:** These authors contributed equally (Cameron Myhrvold, Jessica K. Polka).

## Abstract

Subcellular organization is critical for isolating, concentrating, and protecting biological activities. Natural subcellular organization is often achieved using co-localization of proteins on scaffold molecules, thereby enhancing metabolic fluxes and enabling co-regulation. Synthetic scaffolds extend these benefits to new biological processes, and are typically constructed from proteins or nucleic acids. To expand the range of available building materials, we use a minimal set of components from the lipid-encapsulated bacteriophage Φ6 to form synthetic lipid-containing scaffolds (SLSs) in *E. coli*. Analysis of diffusive behavior by tracking particles in live cells indicates that SLSs are >20 nm in diameter; furthermore, density measurements demonstrate that SLSs contain a mixture of lipids and proteins. The fluorescent proteins mCitrine and mCerulean can be co-localized to SLSs. To test for effects on enzymatic production, we localized two enzymes involved in indigo biosynthesis to SLSs. We observed a scaffold-dependent increase in indigo production, showing that SLSs can enhance metabolic reactions.

## Introduction

Intracellular protein localization is an important mechanism for regulating the function and activity of a variety of cellular processes. Localization influences protein activity by changing the local substrate concentration, access to cofactors and changes in the local environment. To take advantage of this phenomenon, cells frequently organize enzymes into larger protein complexes, such as fatty acid synthetase^1^ and the nonribosomal peptide synthetases and polyketide synthetases of *Bacillus subtilis*^2^. In some cases, these natural scaffolds can involve complex geometry, as found in bacterial microcompartments^3,4^, or consist of a mixture of proteins and nucleic acids, such as the telomerase complex^5^. These complexes can be viewed as scaffolds that co-localize and organize proteins.

Inspired by the diverse set of scaffolds found in nature, synthetic biologists have sought to organize enzymes and other functional molecules on synthetic scaffolds. A pioneering example in 2009 by Dueber *et al.* showed that a protein scaffold could be used to enhance the production of mevalonate^6^. Since then, engineered scaffolds made from proteins or nucleic acids have been effectively used to enhance the activity of various enzymes or signaling proteins to create novel cellular functions^6^–^10^. Both classes of scaffolds have limitations, however: protein scaffolds are limited in spatial complexity, and RNA scaffolds are intrinsically susceptible to enzymatic degradation in the absence of chemical modification^11^. To address some of these limitations, it would be useful to incorporate new types of materials into synthetic scaffolds *in vivo*.

Lipids present a unique and interesting option as a scaffold building material. Lipids can serve as anchors for membrane proteins, and they can also form membrane barriers that allow for selective transport of small molecules in and out of compartments. However, the complexity of membrane-bound organelles preclude their use as a minimal chassis for engineering spatial organization, although existing compartments have been successfully modified^12^ or transplanted to other organisms^13^.

In order to incorporate lipids into synthetic scaffolds, one needs a simple system in which the co-assembly of lipids and proteins of interest can be controlled. A candidate for such a system comes from the bacteriophage Φ6. Unlike most bacteriophages, Φ6 contains a proteinaceous nucleocapsid surrounded by an envelope containing lipids and several membrane proteins^14,15^. During its normal life cycle, Φ6 infects the bacterium *Pseudomonas phaseolicola*^16^ via a membrane fusion event^17^, and thus the membrane is a crucial part of an infectious viral particle. Furthermore, buoyant intermediates in the assembly of Φ6 have been identified, both in the native host during infection^18^ and when a subset of viral proteins are expressed in *Escherichia coli*^19^, consistent with the formation of a lipid-containing structure. More recently, Sarin and coworkers showed that when just 3 viral proteins (P8, P9, and P12) were expressed in *E. coli*, circular particles containing a mixture of lipids and proteins could be purified and imaged with cryoelectron microscopy^20^. Inspired by this work, we endeavored to use Φ6 assembly intermediates as synthetic scaffolds.

Here we report the production of a new class of scaffolds that contain both proteins and lipids. This composition offers the prospects of large size, hydrophobicity, and the incorporation of membrane proteins. We design and build synthetic lipid-containing scaffolds (SLSs) in *E. coli* using parts derived from the bacteriophage Φ6 (Fig. 1A). Specifically, our synthetic system uses two Φ6 proteins: the major membrane protein P9 and a non-structural protein, P12, that is required for particle formation^19^. By fusing various proteins of interest to the C-terminus of P9, we can localize them to our scaffolds. We characterize our scaffolds using fluorescence microscopy and equilibrium flotation centrifugation, and use SLSs to co-localize the fluorescent proteins mCerulean and mCitrine. In addition, we use SLSs to localize two enzymes involved in indigo production, and we demonstrate that the co-localization enhances indigo production. This approach should be generalizable to a wide range of enzymes.

**Figure 1.**
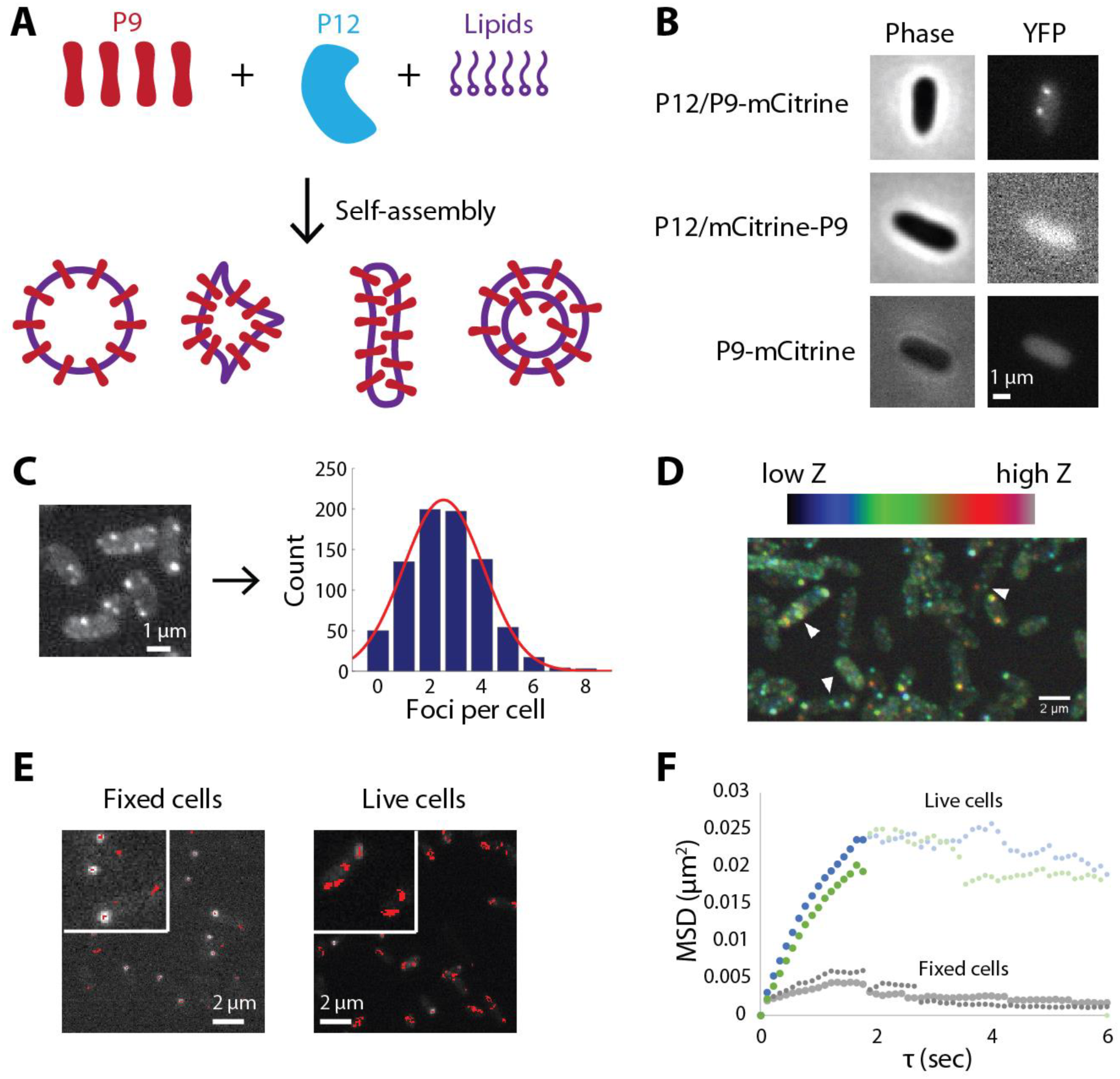
Synthetic lipid scaffolds (SLSs) form discrete foci in *E. coli*. (A) SLSs were designed to self-assemble from the Φ6 phage structural protein P9 in the presence of the required assembly factor P12.^18,19^ (B) C-terminally (but not N-terminally) tagged P9 forms foci in *E. coli*, and P12 is required for this process. (C) The mean number of foci per cell is 2.5 (N = 797 cells). (D) SLSs are present in the middle z slices of a confocal stack. White arrowheads indicate green (mid-Z stack) foci in cells with both blue (low Z) and red (high Z) foci. (E) Particle tracking and subsequent diffusion analysis (F) suggest foci move slower than ribosomes and flagellar basal bodies, suggesting a diameter in excess of 20 nm. Insets in panel E are magnified by a factor of 2.

## Results and Discussion

To directly visualize synthetic lipid scaffolds in *E. coli* cells, we fused the monomeric fluorescent proteins mCerulean and mCitrine to the major Φ6 membrane protein P9. Using fluorescence microscopy, we found that the C-terminal P9 fusion was capable of forming foci in the presence of P12, whereas the N-terminal fusion was not (Fig. 1B). No foci were formed in the absence of P12 (Fig. 1B). While the number of foci per cell varied with the induction level, cells induced for 2 hours using 1 mM arabinose contained a mean of 2.5 foci (Fig. 1C). While foci did not adhere closely to the cell periphery, they were excluded from the central region of the cell, suggesting either transient membrane association or exclusion from the nucleoid (Fig. 1D, for larger images see Supplementary Figure 1).

Particle tracking of these foci revealed that, on average, they diffuse in a manner consistent with globular particles larger than 20 nm. We acquired movies at 9 frames per second and tracked the movement of the particles over time (Fig. 1E). Plots of averaged mean squared displacements (MSDs) of particles over all possible time intervals (τ) in live cells show distinct regimes of diffusive and confined behavior (Fig. 1F). At small values of τ, the MSD plot is roughly linear, indicating diffusive movement. Since we are observing diffusion in two dimensions, we can use the formula MSD = 4Dτ to calculate a diffusion coefficient of 0.00325 to 0.00385 μm^2^/s (R^2^ = 0.92-0.94) for the particles. Thus, SLSs move slower than ribosomes, which diffuse at 0.04 μm^2^/s^21^. SLSs also move slower than flagellar basal bodies, membrane-embedded structures 22nm in diameter that diffuse at 0.005 μm^2^/s^22^. Therefore, regardless of whether SLSs are cytoplasmic or membrane-associated, they likely exceed 20 nm in diameter. The traces are slightly non-linear, however, suggesting that the behavior of P9 particles may be subdiffusive. This could be caused by transient interactions with cellular components such as the cell membrane, nucleoid exclusion, or the physical crowding of the cytoplasm^23^. For larger images of the traces, see Supplementary Figure 1.

Using equilibrium density centrifugation, we find that particles containing C-terminal (but not N-terminal) fluorescently tagged P9 float with a density identical to lipid-containing organelles. We layered the high-speed pellet of clarified extract prepared from cells expressing P9-mCitrine and P12 (Figure 2A) under a sucrose gradient and performed equilibrium density centrifugation. We found a strong peak of fluorescent signal in fractions with a density of approximately 1.17 g/ml (Figure 2B). This density, while higher than that reported for synaptic vesicles (1.05 g/ml), is similar to mitochondria (1.18 g/ml)^24^ and previous density measurements of Φ6 assembly intermediates (1.18 g/ml)^18,19^. Thus, the C-terminal fusion migrates at a density consistent with incorporation into a lipid-containing particle. By contrast, proteins in solution, which have a density of approximately 1.3-1.4 g/ml, would be expected to form a pellet in our gradients^24^. However, the majority of soluble cellular proteins were not loaded onto the sucrose gradient, having been left in the supernatant of a high-speed spin (Materials and Methods). For example, mCitrine-P9 expressed with P12 (Fig. 2C) was left in the soluble fraction (Fig. 2D), suggesting that N-terminal fusions to P9 likely inhibit its ability to properly insert into the membrane. These results are consistent with the cytosolic localization observed by fluorescence microscopy: N-terminal P9 fusions produce a diffuse fluorescent signal, while C-terminal P9 fusions form punctae (Fig. 1B).

**Figure 2.**
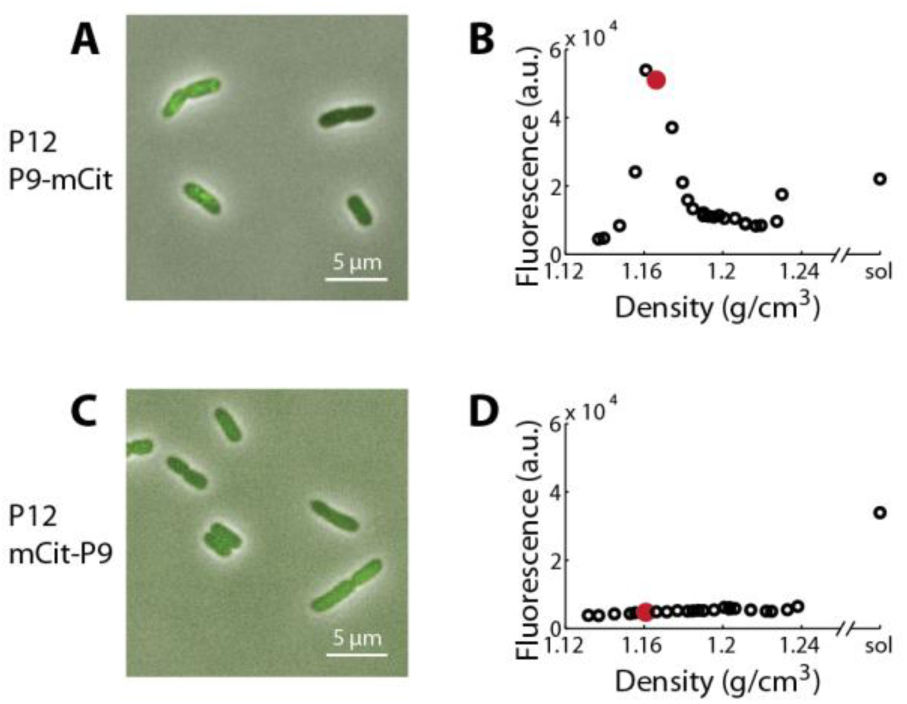
SLSs contain a mixture of lipids and proteins. C-terminally tagged (but not N-terminally tagged) P9 forms puncta in cells (A and C) and floats with a density consistent with lipid-containing particles in an equilibrium density gradient (B and D). Fluorescence measurements were made using a 96-well plate reader (see Methods for details). Fractions highlighted in red in panels B and D were imaged using negative stain TEM (Supplementary Figure 2).

While only the gradient containing C-terminally tagged P9 shows detectable levels of mCitrine, the 1.17 g/ml fractions from both gradients have vesicles that can be observed by negative stain TEM (Supplementary Figure 2A). This is likely because our purification scheme enriches for vesicles in these fractions regardless of the presence of lipid-containing P9 foci (Supplementary Figure 2B); these are likely formed from cell membranes during lysis. The background of these vesicles is too high to reliably detect P9-containing particles by this method. The assignment of any specific morphology to P9 foci is further complicated by the fact that we cannot detect any intracellular particles particular to cells with P9 foci by electron microscopy after high pressure freeze substitution of whole cells (Supplementary Figure 2 C-F). Therefore, we consider SLSs to be discrete, but amorphous lipid-containing particles.

Multiple isoforms of P9 co-localize on synthetic lipid scaffolds. We expressed P12 and P9-mCerulean under control of the arabinose promoter and P9-mCitrine under control of the tet promoter. In the absence of fluorescent fusions, only autofluorescence is observed (Fig. 3A). When only P9-mCitrine is expressed, we observe it distributed throughout the cytoplasm (Fig. 3B). When P12 and P9-mCerulean are expressed, we observe foci in the CFP channel only (Fig. 3C). When P12 and both P9 fusions are expressed, we observe co-localization between mCitrine and mCerulean foci (Fig. 3D, for additional larger images see Supplementary Figure 3). The Pearson’s R value is 0.77, and the ratio of randomized ≥ actual R values is 0.00, suggesting that this correlation is not likely to be due to random overlap (Supplementary file 1). Together, our results indicate that SLSs could be used to scaffold at least two proteins of interest.

**Figure 3.**
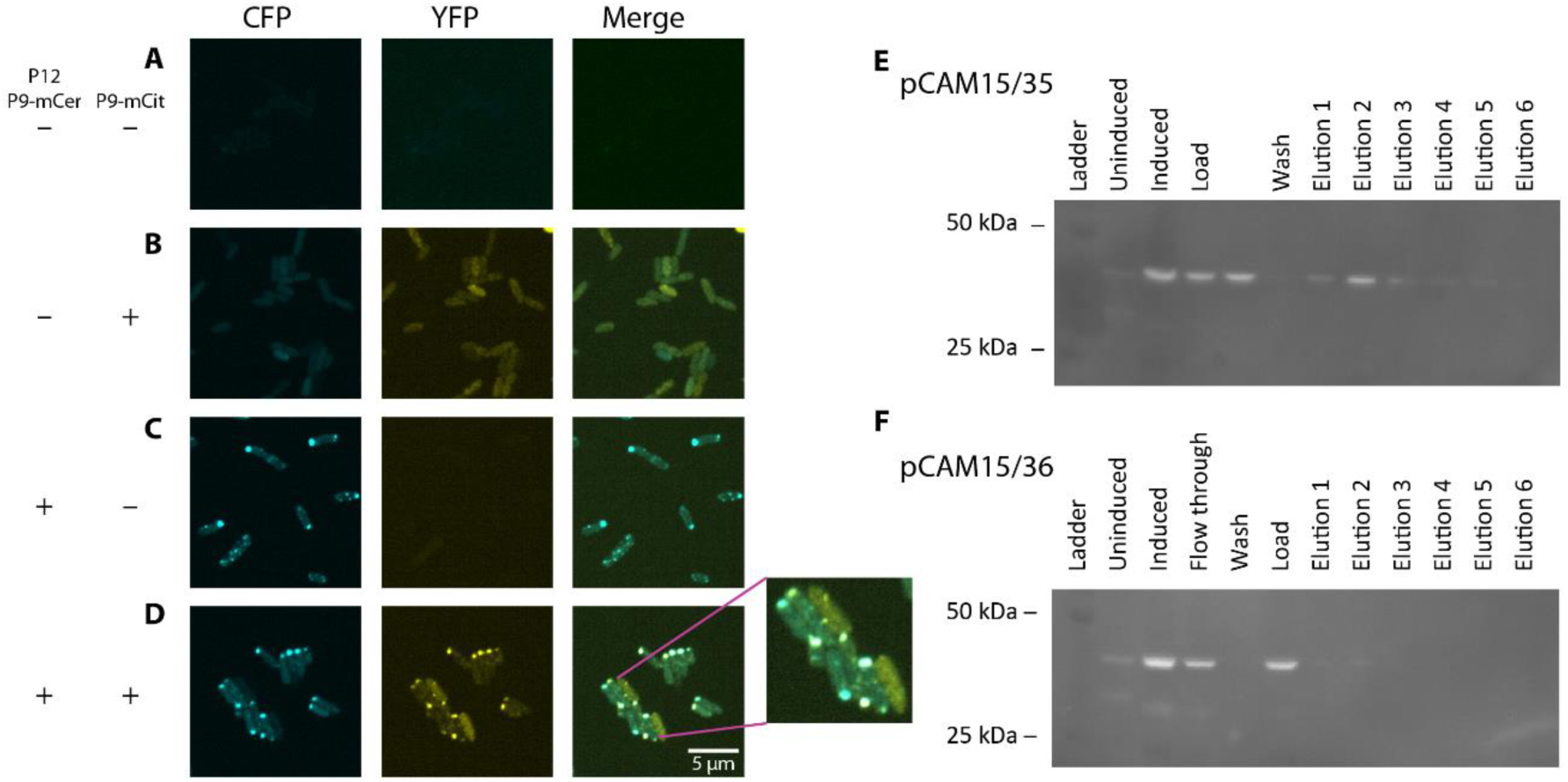
SLSs can co-localize multiple fluorescent proteins. (A) Cells lacking fluorescent P9 fusions are not fluorescent. (B) Cells carrying a plasmid expressing P9-mCerulean are fluorescent, but do not form foci. (C) Cells carrying a second plasmid expressing P12 and P9-mCitrine form foci. (D) In the presence of both plasmids, punctae are visible in both channels. All cells were fixed with 1% (v/v) formaldehyde before imaging. The inset in panel D is magnified by a factor of 2 relative to the other images. (E) P9-mCitrine can be pulled down with P9-His by purification on a nickel column. (F) P9-mCitrine cannot be pulled down by His-P9 under identical conditions.

P9-6His fusions interact with P9 isoforms when co-expressed. We isolated the high-speed insoluble fraction from cells expressing P9-mCitrine and either 6His-P9 or P9-6His and loaded it onto a nickel column. The eluate of the P9-6His preparation contains P9-mCitrine fusions that can be detected by Western blot with anti-GFP antibodies (Fig. 3E). The recovery of P9-mCitrine from a P9-6His affinity purification experiment indicates that these two isoforms of P9 can physically associate with one another. This further implies that at least a fraction of P9 C-termini are exposed to the cytosol, rather than buried within the lipid scaffold, and demonstrates that a protein and an affinity tag can co-localize to SLSs. The eluate from 6His-P9, however, contains very little P9-mCitrine (Fig. 3F), suggesting that the N-terminal 6His fusion, like the N-terminal mCitrine fusion described above, likely does not localize to our scaffolds.

Synthetic lipid-containing scaffolds enhance the production of indigo by co-localizing enzymes. We fused two enzymes involved in indigo biosynthesis, TnaA and FMO, to the C termini of two different copies of P9 and expressed them in the presence or absence of P12 (Fig. 4A). We quantified the resulting indigo production levels of these two strains using a standard curve (Supplementary Figure 4A) and found that strains expressing P12 produced 2-3-fold more indigo than strains that did not express P12 (Fig. 4B). Strains with transmembrane domain deletions in P9 that prevent localization to the particles (P9ΔTM, Supplementary Figure 4B, Supplementary Table 1) and strains that did not express one of the enzymes did not produce detectible levels of indigo. This change in indigo production was not due to a change in the levels of P9-TnaA or P9-FMO, as measured by Western blotting (Fig. 4C, 4D). Thus, we conclude that SLSs can enhance the production of indigo. In the future, SLSs could be used to enhance the production of other metabolites of interest, such as drugs, fuels, or plastics. This new type of scaffold may provide special advantages for processes that involve hydrophobic intermediates or products.

**Figure 4.**
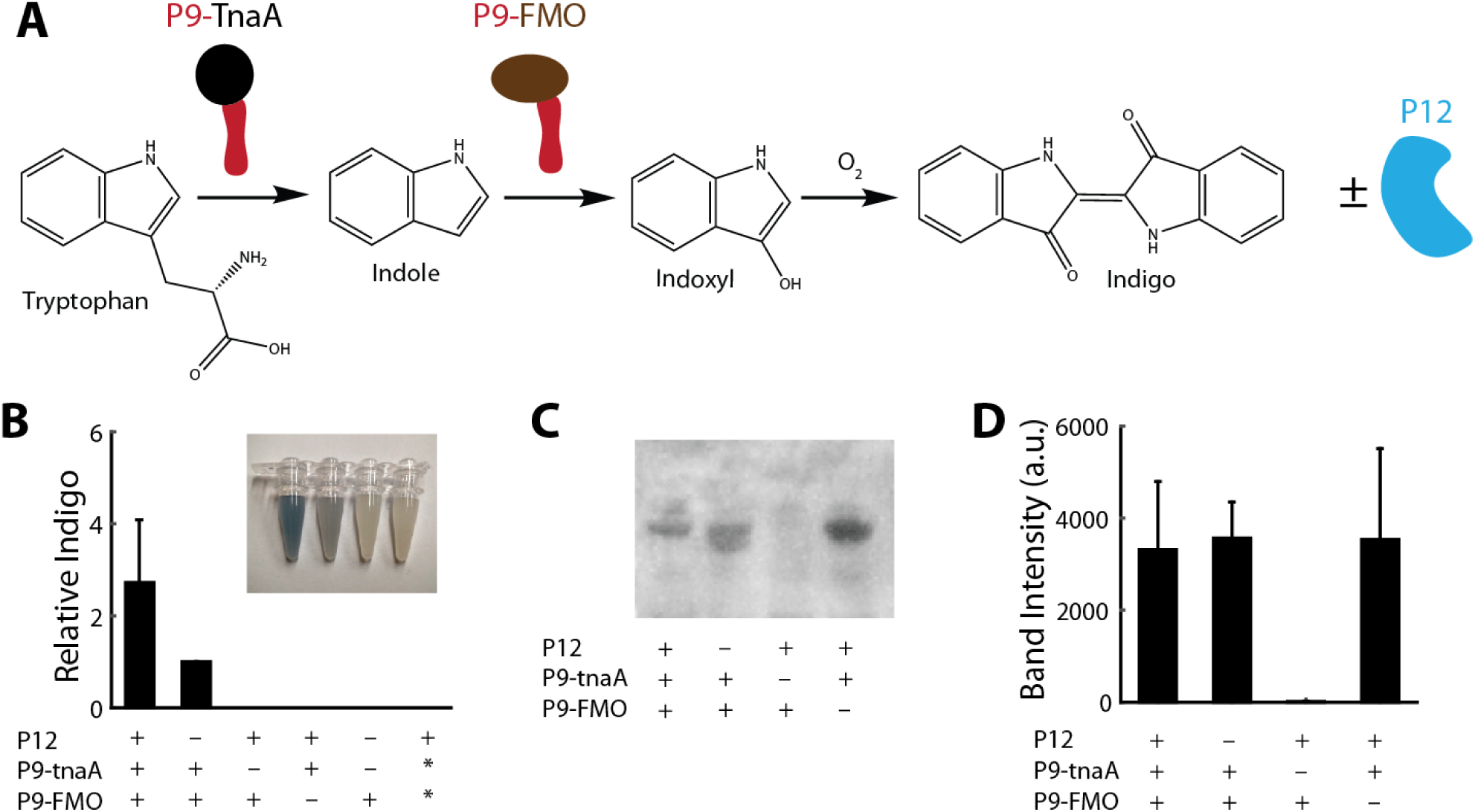
Localization to SLSs enhances the biosynthesis of indigo. (A) A schematic of the proteins expressed for indigo production is shown, along with chemical structures of the relevant metabolites. (B) Indigo production values are shown for strains expressing P9-6His-TnaA and P9-FLAG-FMO fusions in the presence or absence of P12. Data are normalized such that production values for strains without P12 are equal to 1. Error bars indicate one standard deviation based on N=4 experiments. Asterisks indicate mutant forms of P9 that do not contain the transmembrane domain (P9ΔTM). Indigo production was insufficient in the rightmost 4 samples to allow for quantification. (C) Expression of P9-6His-TnaA and P9-FLAG-FMO was measured using Western blotting. (D) Band intensities were quantified to compare expression in the presence and absence of P12 (D). Error bars indicate one standard deviation based on N=3 experiments.

## Methods

### Strains and culturing

For a list of strains and plasmids used in this study, see Supplementary Table 1. For plasmid maintenance, ampicillin was used at a final concentration of 100 μg/ml, carbenicillin was used at a final concentration of 100 μg/ml, and spectinomycin was used at a final concentration of 100 μg/ml. Unless indicated otherwise, cells were grown in LB media. Cells were induced to express proteins with up to 1mM arabinose and 200 ng/ml anhydrous tetracycline (aTc) for 3 hours prior to imaging or purification. Plasmids were constructed using PCR with Q5 or Phusion polymerase (New England Biolabs) and Gibson assembly. Primer sequences are listed in Supplementary Table 2. The P9ΔTM mutant was constructed by site-directed mutagenesis to delete the entire transmembrane helix of P9. The transmembrane helix of P9 was identified using the transmembrane helix prediction server TMHMM^25^, and site-directed mutagenesis was performed using the Q5 site-directed mutagenesis kit (New England Biolabs). Primers are listed in Supplementary Table 2. QIAprep spin or QuickLyse kits (Qiagen) were used for isolating DNA, and DNA Clean & Concentrator™-5 kits (Zymo Research) were used for PCR purification. Plasmid inserts were confirmed by Sanger sequencing (Eton Bioscience or Genewiz, Inc.).

### Particle purification

Cells from a 500 ml culture were harvested by centrifugation and resuspended in 20 ml lysis buffer (containing 200 mM KCl, 10 mM Tris HCl at pH 7.5, 1 mM DTT, and 1 mM MgCl_2_). Immediately after addition of 1 mM phenylmethylsulfonyl fluoride (PMSF), cells were passed through a French Press at approximately 14,400 psi. Lysates were clarified by spinning for 20 minutes at 4,000 rpm at 4 °C in tabletop centrifuge; membranes were isolated from this supernatant by spinning for 110 minutes at 80,000× *g* at 4 °C in a MLA 80 rotor. The insoluble pellet from this spin was resuspended in a small volume (2-5 ml) of lysis buffer containing 62% (w/v) sucrose, and 2 ml of this was layered atop a 1 ml 67% w/v sucrose cushion, in a modification of a protocol described by Laurinavičius *et al.*^26^. Subsequently, 7 ml of 57%, 6ml of 48%, 6ml 39%, and 2 ml 30% sucrose were added. All sucrose fractions were prepared w/v in lysis buffer. Gradients were spun at 32,000× *g* at 4 °C for 86 hours. Fractions were taken for negative stain EM, SDS-PAGE, and fluorimetry by plate reader (Perkin Elmer Victor 3 1420 Multilabel Plate Reader).

### Time lapse microscopy

Cells were either fixed in 1% formaldehyde in PBS or left untouched. They were applied to a 2% agarose pad prepared with M9 + 1% glucose and imaged on a Nikon TE 2000 inverted epifluorescence microscope equipped with a Lumencor LED fluorescence illuminator an Orca ER CCD camera. Using a burst acquisition mode in Nikon Elements AR, images were captured at the maximum frame rate of the camera (8.95 frames/second). Particles were detected and tracked in these movies with u-track^27^ and mean squared displacement analysis was performed using MATLAB.

### Co-localization analysis

Analysis was performed with FIJI^28^. Source images were created by summing 10 individual images of the same field of cells collected from fixed samples. These images were background-subtracted (rolling ball radius of 50 pixels) and both channels were averaged, smoothed (Gaussian blur radius of 2 pixels), and thresholded to create a mask of the cells. This mask, along with the background-subtracted original images, served as inputs for FIJI’s “coloc 2” plugin with a PSF size of 3 pixels and 100 iterations. The output of the analysis is supplied as a supplemental file.

### Electron microscopy

For negative staining, 4 μl of sample was applied to a glow-discharged carbon-coated 200 mesh formvar copper grid (Electron Microscopy Sciences). After 30 seconds, the sample was wicked off with filter paper, washed briefly 3 times in lysis buffer (see above), and 3 times in 0.75% uranyl formate. For sections, cells were subjected to high-pressure freeze substitution followed by Epon resin embedding and staining with uranyl acetate and lead citrate by the Conventional Electron Microscopy Facility at Harvard Medical School. All grids were visualized with a Tecnai G2 Spirit BioTWIN Electron Microscope.

### Western blotting

Samples were run on a 12% Bis-Tris gel in MES buffer. After transfer to nitrocellulose, blots were incubated for 1 h in 2% BSA and/or 5% milk, and then 1.5 h in antibody (Abcam AB1187) rabbit pAb to 6X His (HRP conjugated) at 1:5000 or mouse mAb to GFP (Roche 11814460001) at 1:1000. Chemiluminescence was imaged after Western Lightning ECL treatment. Band intensities were quantified with the “Gels” feature of FIJI.

### Confocal microscopy

Cells were imaged using a Nikon Ti motorized inverted microscope equipped with 100x Plan Apo NA 1.4 objective lens, combined with a Yokagawa CSU-X1 spinning disk confocal with Spectral Applied Research Aurora Borealis modification. For CFP imaging, an 80 mW 445 nm solid state laser with a triple pass dichroic mirror (Chroma) and a 480/40 emission filter (Chroma #849) was used. For YFP imaging, a 100 mW 515 nm solid state laser with a triple pass dichroic mirror (Chroma) and a 535/30 filter (Chroma # 1121) was used. Imaging was performed with a Hamamatsu ORCA-AG cooled CCD camera. Metamorph software was used for image acquisition. For z-stacks, seven optical sections with a spacing of 0.5 microns were acquired, and the data are displayed as maximum z-projections. Brightness and contrast were adjusted uniformly using ImageJ version 1.48 or FIJI.

### Indigo production

For indigo production experiments, cells were cultured overnight in LB supplemented with carbenicillin (100 μg/ml final concentration) and spectinomycin (100 μg/ml final concentration) and incubated at 37 °C with shaking. Overnight cultures were diluted 10^−5^, 3x10^−6^, and 10^−6^ and plated on agar plates containing LB supplemented with carbenicillin and spectinomycin either with inducers (1 mM arabinose and 100 ng/ml anhydrous tetracycline) or without inducers. Plates were incubated overnight at 37 °C, then at room temperature for 6-7 days before harvesting.

Cells were harvested by scraping all of the cells from a petri dish into 1 ml of milliQ H_2_O in a 1.5 ml microcentrifuge tube, and vortexing thoroughly to resuspend the cells. The cells were then pelleted by centrifugation for 10 minutes at maximum speed (~21,000 × g) in a table-top microcentrifuge. The supernatant was removed, and cells were resupsended in 1 ml of milliQ H_2_O. Some samples were split in half so that protein expression levels could be measured by Western blotting. Samples were then pelleted a second time, and resuspended in an equal volume of dimethylformamide (DMF) for lysis. To ensure complete lysis, samples were incubated at 37 °C for 8-16 hours with shaking in the dark.

After lysis, cell debris was pelleted by centrifugation for 10 minutes at maximum speed (~21,000 × g) in a table-top microcentrifuge. 100 μl of supernatant from each sample was transferred to wells of a 96-well plate. Indigo concentrations were measured based on the absorbance at 600 nm on a 96-well plate reader (Perkin Elmer Victor 3 1420 Multilabel Plate Reader). We used polypropylene plates from Eppendorf (catalog # 951040005), as they are not soluble in DMF. Absorbance readings were compared to a standard curve (see Supplementary Figure 4A for an example of a standard curve) based on 2-fold serial dilutions of indigo (purchased from Sigma) in DMF to calculate indigo concentrations.

Cell density counts were obtained using a MACSquant VYB flow cytometer (Miltenyi Biotec). Briefly, washed cells were serially diluted by a factor of 10^−3^ in 1× PBS, and cell densities measured using the MACSquant by collecting 100,000 events per sample and gating based on forward and side scatter. We calculated the indigo produced in milligrams per 10^9^ cells based on these cell density measurements. Indigo concentrations were normalized relative to the unscaffolded samples to quantify the effect of scaffolding on indigo production.

## Acknowledgements

We thank Minna Poranen and Leon Mindich for providing Φ6 expression plasmids, and thank Leon Liu and Tobias Giessen for discussions. We are grateful to the Nikon Imaging Center at Harvard Medical School and to Maria Ericsson and Elizabeth Benecchi at the Harvard Medical School Electron Microscopy facility. This work was supported by the Defense Advanced Research Projects Agency Living Foundries grant HR0011-12-C-0061 and the Wyss Institute for Biologically Inspired Engineering. CM. was funded by the Fannie and John Hertz Foundation, J.K.P was funded in part by the Jane Coffin Childs Foundation.

